# Targeted isolation of bacteria with potential to competitively exclude *Staphylococcus aureus* in the upper respiratory tract of pigs

**DOI:** 10.1101/2024.06.11.598526

**Authors:** Abel A. Vlasblom, Shriram Patel, Peadar G. Lawlor, Marcus J. Claesson, Daniel Crespo-Piazuelo, Julia Eckenberger, Chloe E. Huseyin, Christian Elend, Jaap A. Wagenaar, Aldert L. Zomer, Birgitta Duim

## Abstract

Considering global antimicrobial resistance (AMR) prevalence, alternative or complementary strategies to antimicrobial use, are of interest. Livestock-associated methicillin-resistant *Staphylococcus aureus* (LA-MRSA) is of particular interest as despite significant AMU reduction, LA-MRSA prevalence in pig husbandry has not decreased. We performed targeted isolation of bacterial species with potential antagonism against LA-MRSA in pig farms. Duplicate piglet nasal swabs from three European countries (Germany, Ireland and The Netherlands) were taken longitudinally from birth up to 10 weeks, one for amplicon sequencing and qPCR, and the other was cryopreserved for culturing. We identified potential probiotic species by anticorrelation analysis of bacterial abundance from amplicon sequencing data with quantitative *S. aureus* estimates from qPCR data from the samples. A literature-screen was performed on the species identified, to determine their probiotic potential. Following this, 1302 isolates were grown from selected cryopreserved swabs and identified using MALDI-TOF and additional 16S rRNA gene sequencing to isolate the anticorrelating species. Ninety-five isolates of interest were screened for absence of tetracycline resistance and hemolytic activity and whole genome sequencing was conducted to verify their taxonomy and to assess their AMR and virulence gene profile. Additional phenotypic antimicrobial resistance testing selected three different *Lactococcus lactis* strains. During an *in vitro* challenge using spent medium, all three strains demonstrated inhibition against two *S. aureus* strains. These *L. lactis* strains may have the ability to be used safely to reduce LA-MRSA carriage in the nasal passages of pigs but further *in vivo* testing is necessary to confirm this potential.

**Importance:** An approach to tackle antimicrobial resistance in livestock is to use competitive exclusion agents. We employed a sequencing guided isolation approach to identify bacterial species with *in silico* antagonism to livestock-associated *Staphylococcus aureus* (LA-MRSA) in the nasal microbiome of piglets. Based on this, three *Lactococcus lactis* isolates were found to be suitable for further probiotic testing. This strategy can be used to effectively identify, isolate, and screen bacteria that live in antagonism with opportunistic pathogens such as LA-MRSA in a complex microbiome.

## Introduction

Livestock makes up 62% of the global mammal biomass (1, 2). Factors in livestock husbandry such as dense housing, poor welfare, poor environmental conditions, and inadequate biosecurity, expose and predispose animals to (bacterial) infections (3–5). Historically, antimicrobials were liberally used to treat and prevent bacterial infections in animal herds (6). In light of increased global antimicrobial resistance (AMR), prudent antimicrobial use (AMU) is now considered essential (7). As a result, significant AMU reduction has been achieved in many European countries (8, 9). In the EU (10, 11) and globally, AMU related policies or stewardship programs are on the agenda and becoming the norm (12, 13). It has been shown that AMU reduction can reduce resistance in indicator species (14–16). However, the reduction of antimicrobial resistance appears to be host and bacterial species dependent. For example, despite significant AMU reduction in the Netherlands, livestock associated methicillin-resistant *Staphylococcus aureus* (LA-MRSA) prevalence in pig production has not decreased (17). Therefore, alternative, or supplementary pathogen reduction strategies, to antimicrobials, would be a valuable tool in livestock husbandry. Alternatives can consist of both preventative and therapeutic measures (18). Here, we aimed to identify (probiotic) bacterial strains, that can be applied as a preventative strategy to reduce LA-MRSA carriage in pigs. We focused on LA-MRSA as it colonizes animals globally on pig farms and is a health risk for humans (19). For example, in 2020, 12% of Dutch human clinical MRSA isolates were identified as LA-MRSA (20). Another example is LA-MRSA making up 35% of the detected MRSA strains in a German hospital in a livestock dense region (21).

The use of probiotic organisms with competitive exclusion properties can benefit host health (22). Their competitive-exclusion mechanisms can include the production of antimicrobial substances, epithelial barrier function enhancement, immunomodulation, or signaling via the nervous system (23). Such a relationship between microbes in the nasal cavity and *S. aureus* carriage has been suggested in humans (24–29) and in pigs (30–32). Competition between nasal species, for example between a probiotic and *S. aureus*, could occur through competition for adhesion sites and nutrients, or by antibiosis or induction of host defenses (33). In a complex microbiome, these mechanisms can result in a negative association between the probiotic and target pathogen. In this study, we attempted to identify these associations in a longitudinal piglet nasal-swab dataset through anticorrelation analysis (Vlasblom *et al.*, 2024 (34)). We specifically aimed to screen and identify potential probiotic species in the piglet nasal cavity that exclude *S. aureus*, by anticorrelation analysis of bacterial species abundance from amplicon sequencing data with quantitative *S. aureus* estimates from qPCR data. We furthermore aimed to assess, following international guidelines, the suitability of these bacterial species for use in animals both *in silico* and *in vitro*.

## Results

### General study outline

In a previous study by Vlasblom et al., 2024 duplicate nasal swabs from healthy pigs were longitudinally collected, in 9 farms in Germany, Ireland, and the Netherlands. DNA was extracted from one swab to investigate the sample bacterial composition through 16S rRNA and *tuf* gene amplicon sequencing, and the second swab was cryopreserved at -80°C for isolation of bacteria (34). Here, *in-silico* analysis of longitudinal nasal swab data combined with *S. aureus* specific qPCR data was conducted to identify species having a negative association with *S. aureus*. Species having the highest negative correlation were subsequently targeted in an isolation effort from nasal swab material. Isolates obtained were identified using MALDI-TOF and 16S rRNA sequencing and screened bioinformatically and using *in vitro* assays for ‘safety’, meaning that these species were not considered pathogenic and had a low-risk status as defined by several regulatory organisations. The ESFA guide on the characterisation of microorganisms used as feed additives or as production organisms, was used to exclude any species with potential virulence or antimicrobial resistances. A workflow summary can be found in Figure 1.

**Figure 1.**
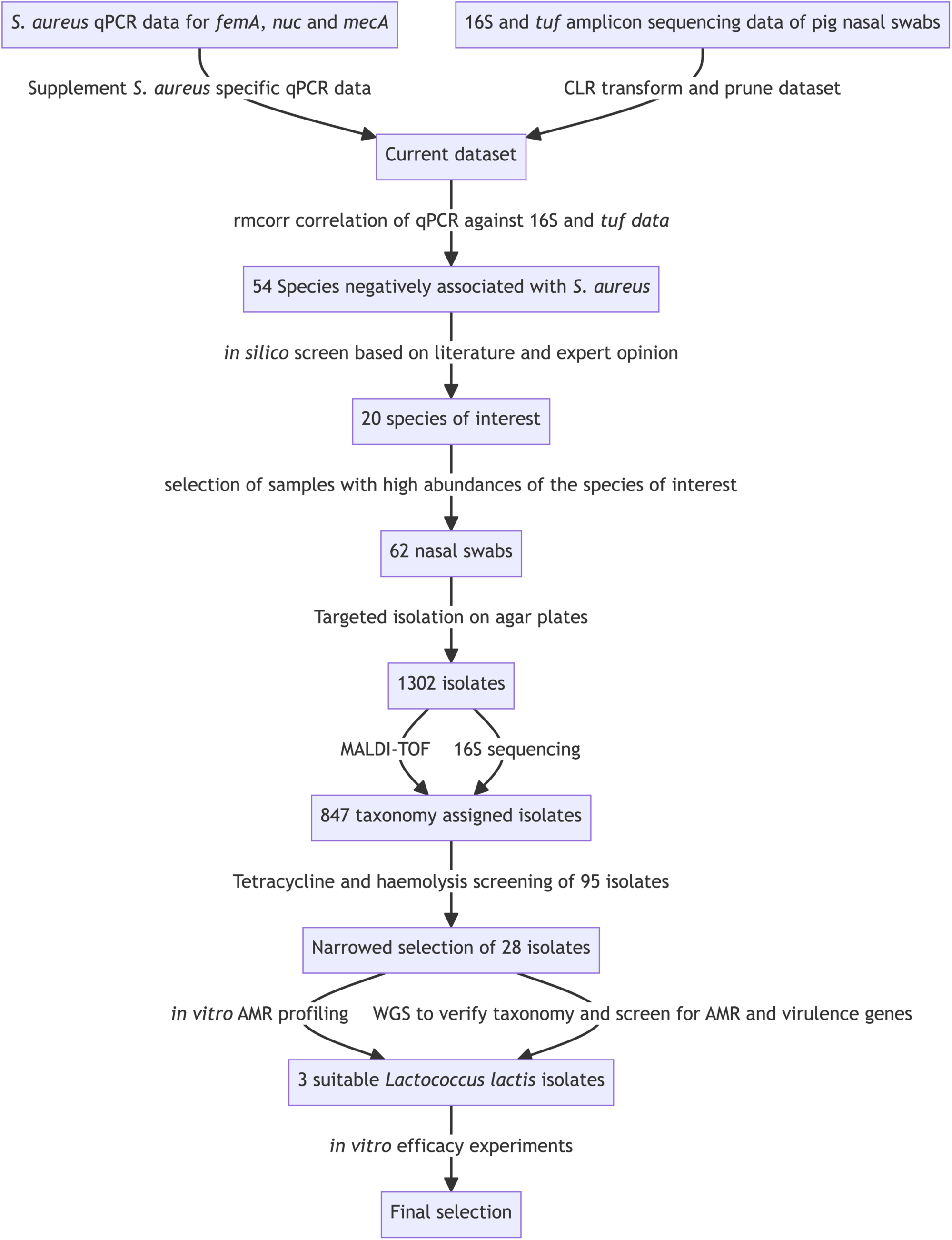
Workflow of the study

### Identification of species negatively associated with *S. aureus* by enriching amplicon sequencing data with species specific qPCR-data

Longitudinal amplicon sequencing data (based on the V3-V4 16S rRNA and *tuf* genes) from previous work (34) was correlated with *S. aureus* specific qPCR data (*femA* and *nuc* genes), to identify species negatively linked with *S. aureus*. qPCR supplementation was chosen due to its increased *S. aureus* detection compared with amplicon sequencing on its own, likely due to low *S. aureus* load and the sensitive targeted amplification of qPCR. *S. aureus* qPCR estimates were correlated with centered log-ratio (CLR) abundances of 16S rRNA and *tuf* gene amplicon sequencing taxa using the R-package rmcorr. We chose rmcorr to account for repeated measurements in our longitudinal dataset. This analysis yielded 54 unique species from 34 genera, that had a negative correlation with *S. aureus*, with an r value < -0.1 and a p-value < 0.1 over the *tuf* and 16S rDNA data. We found around 15 negative species associations with *femA* or *nuc* per country in the *tuf* data. While in the 16S rDNA data, the Netherlands were overrepresented with 47 and Germany was underrepresented with six anticorrelations identified (details can be found in supplementary Table 1).

### Literature review and expert screen identified species of interest

All species found to be negatively associated with *S. aureus* were reviewed in literature for probiotic suitability. A summary of this screen is found in Supplementary Table 1. In the literature screen we paid particular attention to removal of (potential) pathogens to pigs and other vertebrates. The parameters, chosen to exclude pathogens from the potential probiotic candidates narrowed the list down to species that were either rarely/under-described or had a low-risk status as defined by the GRAS list in the US or QPS list from EFSA or the German BAuA/ABAS’s “Technical Rules for Biological Agents” list (TRBA). This screen narrowed our selection to 20 bacterial species of interest from 11 genera (Table 2). The species *Carnobacterium inhibens* and *Lactococcus lactis* were identified in both the *tuf* and 16S datasets, albeit in different countries. The strongest average anticorrelation against *S. aureus* was found in the Irish *tuf* dataset by *Jeotgalicoccus marinus*, a species member of the *Staphylococcacae* family that was first described in sea urchins (35). In the 16S dataset the strongest average anticorrelation belonged to *Moraxella boevrei*, a relatively unknown *Moraxella* species first described in 1997 (36).

**Table 1.** The number of anticorrelating species with *S. aureus* (qPCR) for each dataset (*tuf* or 16S) per country with an r < -0.1 and a p <0.1.

| Country | Number of species negatively associated with *S. aureus* |  |
| --- | --- | --- |
|  | 16S | *tuf* |
| The Netherlands | 47 | 15 |
| Germany | 6 | 16 |
| Ireland | 14 | 15 |

**Table 2.** The species of interest as determined by literature screen, and their average negative R-value per dataset and country. The R-value shows the average strength of the anticorrelation in each country.

| Species | Germany | Ireland | the Netherlands |
| --- | --- | --- | --- |
| **16S** |  |  |  |
| *Bergeyella porcorum* |  | -0.11 | -0.18 |
| *Carnobacterium inhibens* | -0.1 |  |  |
| *Chryseobacterium taklimakanense* |  | -0.14 | -0.14 |
| *Lactobacillus amylovorus* |  | -0.17 |  |
| *Lactobacillus mucosae* |  |  | -0.12 |
| *Lactobacillus salivarius* |  |  | -0.2 |
| *Lactococcus lactis* | -0.12 |  |  |
| *Lactococcus raffinolactis* | -0.1 |  |  |
| *Leuconostoc pseudomesenteroides* |  |  | -0.14 |
| *Moraxella boevrei* |  | -0.16 | -0.22 |
| *Thiovirga sulfuroxydans* |  |  | -0.13 |
| *Weissella cibaria* | -0.11 |  |  |
| *Weissella viridescens* |  |  | -0.19 |
| *Zoogloea ramigera* |  |  | -0.14 |
| **tuf** |  |  |  |
| *Blautia hansenii* |  |  | -0.17 |
| *Carnobacterium inhibens* |  | -0.24 | -0.14 |
| *Jeotgalicoccus marinus* | -0.16 | -0.27 | -0.23 |
| *Lactobacillus delbrueckii* | -0.15 |  |  |
| *Lactobacillus helveticus* | -0.12 | -0.22 | -0.15 |
| *Lactococcus lactis* |  | -0.25 | -0.18 |
| *Staphylococcus cohnii* |  | -0.19 | -0.19 |
| *Staphylococcus xylosus* |  |  | -0.18 |

### Directed species isolation

To isolate the species of interest, a directed culturing effort was undertaken. The top five samples containing the highest relative abundances (for the *tuf* and 16S amplicon data) of the species of interest were selected. During the literature screening, scientific publications and public websites were reviewed to find suitable growth conditions for each species of interest to facilitate their culturing and isolation (Table 3 and Supplementary Table 1). Species isolation was performed by streaking cryopreserved material of 62 swabs from Germany, Ireland, and the Netherlands on agar plates. When colonies had formed on the agar plates, isolates were sub-cultured from colonies with distinct morphologies per growth condition (28 conditions) to obtain pure cultures which were then characterized with MALDI-TOF analysis (Table 3).

**Table 3.** Growth conditions and the associated number of MALDI-TOF confirmed species.

| Gas mixture | Temp | Agar Medium | Isolates picked | Target genus |
| --- | --- | --- | --- | --- |
| 5% CO2 | 24 | MRS | 19 | *Weissella* |
|  | 30 |  | 217 | *Lactobacillus, Lactococcus, Leuconostoc, Weissella* |
|  | 45 |  | 131 | *Lactobacillus, Lactococcus* |
| Anaerobic | 30 |  | 74 | *Lactobacillus, Lactococcus* |
|  | 45 |  | 64 | *Lactobacillus, Lactococcus* |
| Microaerobic | 37 |  | 8 | *Lactococcus* |
|  | 45 |  | 19 | *Lactobacillus* |
| 5% CO2 | 30 | BHI | 17 | *Blautia, Lactococcus, Staphylococcus* |
|  | 37 |  | 9 | *Blautia* |
| Microaerobic | 37 |  | 10 | *Blautia* |
| 5% CO2 | 30 | Blood | 108 | *Bergeyella, Lactococcus, Moraxella, Staphylococcus* |
|  | 37 |  | 17 | *Bergeyella* |
| Aerobic | 37 |  | 40 | *Lactobacillus, Moraxella, Staphylococcus* |
| Anaerobic | 30 |  | 30 | *Lactococcus* |
| Microaerobic | 24 |  | 5 | *Moraxella, Staphylococcus* |
|  | 37 |  | 26 | *Moraxella, Staphylococcus* |
| Aerobic | 37 | Choc | 5 | *Moraxella* |
| Microaerobic | 37 |  | 9 | *Moraxella* |
|  | 24 |  | 1 | *Moraxella* |
| 5% CO2 | 30 | MA5 | 40 | *Jeotgalicoccus* |
|  | 37 |  | 20 | *Jeotgalicoccus* |
| Anaerobic | 30 |  | 28 | *Jeotgalicoccus* |
| 5% CO2 | 30 | TSA | 34 | *Carnobacterium, Chryseobacterium* |
| Aerobic | 37 |  | 40 | *Staphylococcus* |
| 5% CO2 | 30 | LB | 16 | *Staphylococcus, Zoogloea* |
| Anaerobic | 30 | GYP | 286 | *Carnobacterium, Lactobacillus, Lactococcus* |
| Anaerobic | 30 | NA | 18 | *Chryseobacterium* |
| Microaerobic | 30 | MSA | 11 | *Jeotgalicoccus* |
| **Total** |  |  | **1302** |  |

Brain heart infusion agar (BHI), Columbia Agar with 5% Sheep Blood (Blood), Chocolate Agar (Choc), Glucose Yeast Peptone Agar (GYP), Luria-Bertani Agar (LB), Marine Agar with 5% NaCl (MA5), De Man–Rogosa–Sharpe agar (MRS), Mannitol Salt Agar (MSA), Nutrient Agar (NA), and Trypticase Soy Agar (TSA).

### Species characterization with MALDI-TOF and 16S sequencing

Of the 1302 isolates, grown from the swabs, originating from Germany (475 isolates), Ireland (254 isolates), and the Netherlands (573 isolates), MALDI-TOF assigned taxonomy to 773 isolates (Supplementary Table 2). Eighty-four isolates that could not be identified by MALDI-TOF were 16S rDNA sequenced. Ultimately, we assigned taxonomy to 847 isolates (either at genus or species level) (Supplementary Table 2).

### Tetracycline MIC and haemolysis screen

Of the 847 total isolates that were assigned a taxonomy, 95 isolates belonged to the genera of interest. These isolates were therefore selected for a tetracycline phenotypic resistance test. In this test isolates were spotted on agar plates with increasing tetracycline concentrations. Concentrations were based on the 2018 EFSA guidance “Guidance on the characterization of microorganisms used as feed additives or as production organisms (Table 2)” (37), as the guidance dictates the tetracycline MIC per bacterial species. Only 28/95 isolates passed the tetracycline MIC cut-off values as defined for the assigned genera (Table 4). The 27 isolates passing the tetracycline screen were re-evaluated on their assigned taxonomy by MALDI-TOF or 16S rDNA sequencing, this removed five species with conflicting MALDI-TOF results and reduced the selection to 23 isolates (Table 4). These isolates were subsequently grown on blood agar to assess their haemolytic activity as this is an unwanted trait for a potential probiotic (38). Six isolates showed haemolysis and this reduced the selection to 17 isolates (Table 4).

**Table 4.** Summary of the number of isolates by genus passing the required criteria.

| Assigned genus | No tetR | MALDI-TOF species ID | No alpha or beta haemolysis | WGS taxonomy | No AMR or virulence genes |
| --- | --- | --- | --- | --- | --- |
| Jeotgalicoccus | 2 | 2 | 2 | 0 | 0 |
| Lactobacillus | 6 | 5 | 3 | 1 | 0 |
| Lactococcus | 3 | 3 | 3 | 3 | 3 |
| Leuconostoc | 4 | 2 | 1 | 0 | 0 |
| Moraxella | 6 | 6 | 5 | 0 | 0 |
| Staphylococcus | 4 | 3 | 3 | 3 | 0 |
| Weissella | 2 | 1 | 0 | 0 | 0 |
| **Total** | **27** | **23** | **17** | **7** | **3** |

Each column represents the number of isolates per genus matching the required criteria: phenotypic tetracycline resistance (tetR), species confirmed by MALDI-TOF, absence of alpha or beta haemolysis, and their taxonomy with absence of antimicrobial resistance (AMR) and virulence factors determined by whole genome sequencing (WGS).

### Whole genome sequencing to confirm taxonomy and screen for AMR and virulence genes

The whole genome of the remaining 17 isolates was sequenced to verify their taxonomy and to determine if AMR genes or virulence factors were present. The assigned taxonomy of seven assemblies matched the anticorrelating “species of interest” using the Epi2me, kmerfinder, and TYGS databases (Table 4). Another eight isolates had inconsistent taxonomy assignments, and two isolates belonged to species not selected as a “species of interest”. Next, we identified in the assemblies of the seven matching species the presence of resistance genes, using Resfinder (Table 4). In only three isolates were no AMR genes present. These were *Lactococcus* lactis isolates, which had passed all the required selection criteria. *L. lactis* has GRAS status from the FDA in the US (39) and QPS status within the EU as defined by EFSA (40). These statuses make it favorable for probiotic use. An overview of all performed assays can be found in Supplementary Table 3.

### In vitro AMR profiling of L. lactis

The Minimal Inhibitory Concentration (MIC) for each antibiotic on the panel was determined for the three isolates following EFSA guidance to quantify phenotypic resistance. All three isolates met the MIC cut-offs as determined for *L. lactis* by EFSA (Table 5).

**Table 5.** Characteristics of the selected *L. lactis* isolates.

| Isolate ID | Synonym | Country | Minimal inhibitory concentration (MIC) in µg/mL |  |  |  |  |  |  |  |  |
| --- | --- | --- | --- | --- | --- | --- | --- | --- | --- | --- | --- |
|  |  |  | AMP | CLI | GEN | KAN | CMP | ERY | STR | TET | VAN |
| **IV-F-45-10** | EW-01 | NLD | ≤2 | <=0.5 | 1 | <16 | ≤8 | ≤0.25 | 8 | ≤2 | ≤0.25 |
| **VI-AI-3** | EW-05 | GER | ≤1 | <=0.5 | ≤0.5 | <16 | ≤8 | ≤0.25 | 6 | ≤2 | ≤0.25 |
| **VII-E-8** | EW-04 | NLD | ≤1 | <=0.5 | 1 | <16 | ≤8 | ≤0.25 | 6 | ≤2 | ≤0.25 |

ampicillin (AMP), clindamycin (CLI), gentamicin (GEN), kanamycin (KAN), chloramphenicol (CMP), erythromycin (ERY), streptomycin (STR), tetracycline (TET), vancomycin (VAN)

### In vitro competition of L. lactis with S. aureus

To assess if the three *L. lactis* isolates showed *in vitro* effects on *S. aureus* growth, a spent medium based assay was performed. Two strains of *S. aureus* ATCC 25913 and ATCC 25923 were grown on spent medium (filter sterilized waste medium) made from the cultivated Lactococci. To determine if an inhibiting effect was the result of acidification or nutrient competition, we included three conditions where we either adjusted the pH of the medium to 7.4 (Supplementary Table 4), we added 50% of fresh medium, or we did both. Cultures were grown overnight at 37°C and optical density at 600nm (OD600) was measured every 10 minutes during. From the resulting growth curves, we calculated the maximum growth rate in OD600 per hour and reported this as fraction of growth on fresh BHI (positive control) (Figure 2). The growth rate of ATCC 25913 was almost halved by its own spent medium and by the medium of the three Lactococci. The growth speed of ATCC 25923 was drastically reduced by the Lactococcus-medium, to speeds as low as 5% of the positive control, and to 36% speed by its own spent medium. Adjusting the pH to 7.4 seemed to recover the growth speed of ATCC 25923 to ∼50%, where the growth speed of ATCC 25913 did not increase. For ATCC 25913 we observed that both the addition of 50% BHI, and addition of 50% BHI and adjusted pH, seem to recover the growth speed similarly. This suggests that ATCC 25913 growth speed inhibition was more nutrient dependent, whereas the reduced growth speed of ATCC 25923 was both nutrient and pH dependent. Ultimately, this was a limited test that does not represent the complexity of the microbe-microbe interactions present in the pig nostril, but it does indicate a direct competition though nutrient competition or acidification.

**Figure 2.**
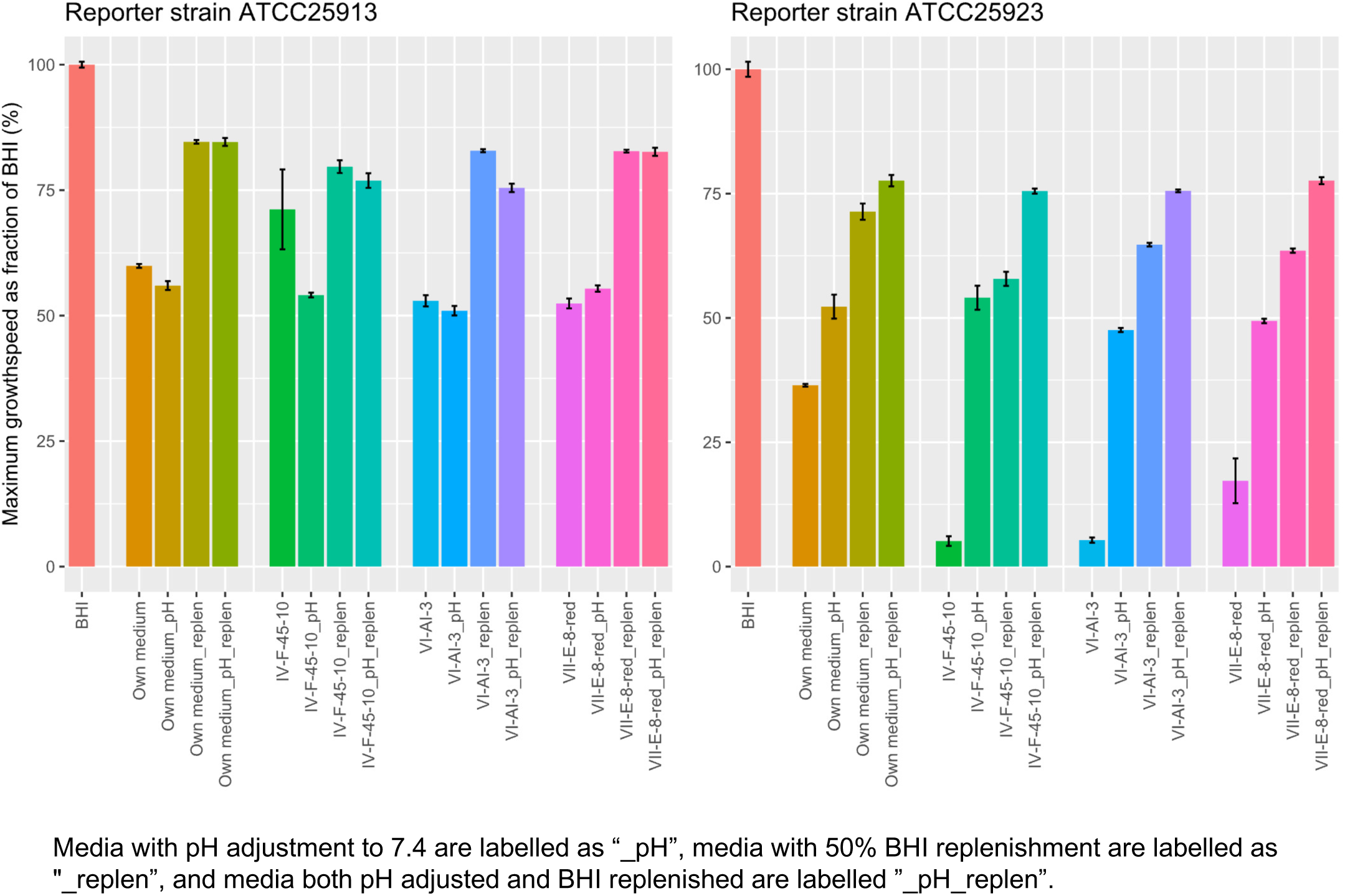
Average growth speed of *S. aureus* ATCC25913 (left) and *S. aureus* ATCC25923 (right) reporter strains grown on spent media from the three *L. lactis* (IV-F-45-10, VI-AI-3, VII-E-8) and on its own spent medium.

## Discussion

### Bacterial species with anti-*S. aureus* associations were observed in pigs from three countries

In this study we identified 54 taxa negatively associated with qPCR-based quantification of *S. aureus* loads, of which we, based on literature review, deemed 20 species of interest for probiotic application. Our earlier study (31) also utilized qPCR to determine *S. aureus* anticorrelating species. However, here we were able to identify taxa down to species level and to identify taxa that were negatively correlated with *S. aureus* in different farms and countries. Furthermore, in the current study we stored duplicate swabs for an isolation effort. Negatively associated species were present in pig nasal samples, often with a weak correlation coefficient. However, these correlations were found in all three countries suggesting robustness. For example, *L. lactis* was found to be negatively associated with *S. aureus* in all three countries (Table 2). This species correlation, although significant, were weak, but the repeated detection of a negative correlation in the data from nine different farms from three different countries suggests a real effect. This effect might be due to the microbiome developing over time with successive nasal colonization by different bacterial species, as our previous work suggested (31, 34). Still, it is important to consider that negative correlations in longitudinal data may be a consequence of confounding effects caused by external factors that shape the nasal microbiome rather than being indicative of direct microbe-microbe interactions (41)

### *L. lactis* as an anti-*S. aureus* probiotic?

Probiotics can be used effectively to decolonize hosts from *S. aureus* as was recently shown with *Bacillus subtilis* in humans (42). The current study resulted in a final selection of three suitable *L. lactis* isolates, meeting guidance by EFSA. The *L. lactis* species is suitable as a probiotic because it has a GRAS and QPS status, is widely used in food production, and is authorized as a European feed additive (39, 40, 43). Furthermore, *L. lactis* has been proposed as a mucosal intranasal vaccination platform (44). Also, the fact that we isolated *L. lactis* from pig samples from the same niche might improve their probiotic potential as this accounts for host and niche specificity complications (45). Apart from the general probiotic potential of *L. lactis*, an *S. aureus* specific inhibiting effect by *L. lactis* in piglet noses is not unthinkable, as these competitive effects have been reported previously. For example, oral *L. lactis* supplementation in a murine model (46), in epithelial cell-line invasion experiments (47) and in fermented cheese (effect was likely pH dependent inhibition) (48). Furthermore, *L. lactis* has been found to change the virulence of *S. aureus* by downregulating its accessory gene regulator *(agr)* system (49). *L. lactis* has also been found to produce *S. aureus* inhibiting biocins (50). Additional inhibiting methods might consist of a quorum-quenching effect, as it has been shown between *Bacillus subtilis* and *S. aureus* in humans (51). Our growth rate assay showed that *S. aureus* reporter strains were inhibited when grown on the Lactococcal spent media. The three *L. lactis* spent media had become acidic (Supplementary Table 4), and the growth rate of ATCC 25923 increased when the spent media was adjusted to pH 7.4. This suggests that the reporter strain was inhibited by acidification, something that *L. lactis* is well known for (52). We also noticed an increase in growth rate of *S. aureus* after nutrient replenishment in the media. Nutrient competition in the nasal cavity has been suggested to be a factor affecting *S. aureus* colonization (29, 33). Based on this study we do not know the exact mechanism by which the *L. lactis* strains might compete with *S. aureus in vivo*, however we do suspect pH and/or nutrient competition are involved. However, other mechanisms such as discussed above, cannot be excluded based on our tests. Their safe status, the earlier observed anti-*S. aureus* effects, absence of relevant AMR (both *in vitro* and *in silico*) and potential virulence determinants (Tables 4 and 5) make these three *L. lactis* isolates interesting candidates for further clinical trials. Additionally, the safety of the isolates for *in vivo* nasal application has recently been established (53) and we plan to investigate their anti-*S. aureus* activity in a randomized controlled field trial.

### A targeted isolation approach resulted in isolation of prospective isolates but leaves room for improvement

Species isolation was performed by selection of nasal swab samples with the highest *in silico* relative abundance per species of interest. This resulted in a targeted isolation effort. This targeted approach limited the quantitative interpretation of the dataset and reduces the potential for systematic description of the culturable species per growth condition, that a systematic culturomics approach (32) would have allowed. However, we knew what species were of interest based on the *in silico* anticorrelation. Therefore, a targeted approach was considered faster and more resource efficient. We submitted 1302 isolates for MALDI-TOF identification, a much lower number of isolates than an untargeted sampling approach would have required.

Knowledge of legislation (e.g. of phenotypic AMR profiles) could inform future study design. For example, by selecting parameters that reduce the number of isolates to screen early on. In this case, we would propose transferring isolates to a 16 mg/L tetracycline containing growth medium after initial colonization on the general growth permitting medium. Because, except for *Lactobacillus plantarum/pentosus* or *Lactobacillus reuteri*, no bacterium is permitted to be used as feed additive or as production organism that grows at this concentration (minimum of 2x above MIC) following EFSA-guidance. Furthermore, tetracycline resistance is widespread in pig farming (54). This could reduce the number of isolates very early on in the effort. Therefore, we suggest that after isolation, isolates are *ab initio* screened for AMR phenotypic resistances (e.g. tetracycline, as it is common) prior to in depth identification efforts. Another early screen could be the determination of haemolysis on blood agar, as several of the species tested produced haemolytic factors, which could limit their applicability as probiotics.

A complication in our workflow arose from needing proper taxonomy assignment. Our workflow started with amplicon sequencing data. However, 16S rDNA amplicon sequencing does not always differentiate species properly (55). We then assigned taxonomy to isolates using MALDI-TOF, but this platform also has been found to have difficulties differentiating closely related species (56). Finally, we assigned isolate taxonomy using WGS. The different methods of assigning taxonomy to the sequenced isolates did not always yield similar results (Supplementary Table 3). An alternative strategy would be to start with shotgun metagenomics instead of amplicon sequencing, for combined detection of functional genes and species taxonomy. However, in an unpublished pilot we noticed that host reads made up around 90-99% of the total nasal swab reads.

### Following legislation might be too stringent for selecting prospective probiotics

Following legislative/guidance documents is essential to select bacterial strains that can be most readily utilized as probiotics within a specific jurisdiction. Therefore, we ultimately selected specific strains that had QPS and/or GRAS status. However, from a fundamental perspective these strains were neither the most abundant nor the most negatively associated with *S. aureus* (Table 2, Supplementary Table 1). When attempting to identify “novel” bacteria for pathogen exclusion strategies, the prior description of an organism might be an overly stringent selection criterium. Many bacteria associated with vertebrate hosts have been found in cases of opportunistic infection. Even for species seemingly suitable for probiotic use, cases of pathogenicity have been described, for example for *L. lactis* in cows (57), and even in humans (58). Even in commercially available probiotics there are many unknowns, and some risks associated with their use. For example commercially available probiotics, including strains of *L. lactis,* have been found to often carry (mobile) AMR genes (59).

We attempted to be lenient in our initial species of interest selection. For example, we included *S. cohnii* as strain of interest, despite previous association with human disease (60). This was based on previous findings that the species has been found to be negatively associated with *S. aureus* in the pig nasal cavity (61). This paired with a favorable safety classification in Germany (62) prompted us to retain the species as a candidate for the purposes of our study. However, ultimately no suitable isolates of this species were obtained after phenotypic and genotypic screening, but when isolates are available of this species that contain the required characteristics, they are of interest for future studies.

Interestingly, organisms that are severely under-described (E.G. *M. boevrei*, and *J. marinus*) showed strong anticorrelation in our data. Again, supporting that in future work, a species should not be excluded from evaluation simply because it does not have for example QPS status (40). Such an approach could result in more effective strains with higher probiotic potential being selected. However, it is important that these strains are subsequently fully safety tested *in vitro* and *in vivo* prior to use. Clinical safety trials could therefore allow strains, already present in the described niche, to be approved for probiotic use, similar to what we have performed for these *L. lactis* strains (53).

### Final remarks

Our study, using a targeted approach, identified three *L. lactis* strains with weak negative correlation with *S. aureus*, suitable for *in vivo* safety and efficacy testing in pigs. The three isolates have subsequently had their safety assessed *in vivo* (53). The three isolates have been found to be safe for nasal administration in pigs. The next step will be to conduct an *in vivo* efficacy study, as *in vitro* assessment of probiotic potential is hindered by the absence of host effects (immunological, spatial) with potential to obscure the effect of the probiotics.

## Materials and methods

### *S. aureus* specific qPCR

DNA extractions from nasal swabs (Vlasblom *et al.*, 2024) were used for qPCR assays. This dataset consisted of nasal swabs that were obtained from 54 piglets born to 27 sows across nine farms equally distributed in Germany, Ireland, and the Netherlands. qPCR assays consisted of *S. aureus* specific *femA* and *nuc* targets in a probe assay, and 16S rDNA qPCR in a SYBR-green assay, using a LightCycler480 (Roche) with primers, reactions and chemistry as described in Rittscher *et al.* (63) Generated *S. aureus* qPCR estimates were normalized between samples using 16S rDNA qPCR bacterial load estimates (31, 63).

### rmcorr anticorrelation of amplicon sequencing data with normalized *S. aureus* specific qPCR data

qPCR results were combined with the *tuf* and 16S count datasets (from the Vlasblom *et al.*,2024) and imported as Phyloseq (64) objects in R. The microbiota count data from *tuf* and 16S datasets were filtered for low abundant taxa with minimum proportion of 0.01 and minimum prevalence of 0.01 using codaseq.filter wrapper from CoDaSeq package (65). The qPCR-data were scaled by total microbial cell counts and log-transformed before correlation analysis. To account for the compositional nature of the microbiota data and to avoid the likelihood of generating spurious correlations, we first imputed the zeros in the filtered abundance data using the count zero multiplicative replacement method (cmultRepl, method = “CZM”) implemented in the zCompositions package (66) and applied a centered log-ratio transformation (CLR) using the codaSeq.clr function in the CoDaSeq package. Associations between species and qPCR data were then obtained using repeated measure correlation analysis from the rmcorr package (67), which determines the relationship between two continuous variables while controlling for between-individual variance. Because we found the highest microbiota variance by country of sample collection (not shown), we performed rmcorr analysis separately for three countries and combined the tables for downstream analysis.

### Literature review of anticorrelating species

Literature review was supported by search-queries in PubMed (https://pubmed.ncbi.nlm.nih.gov/; Aril-July 2021): species-name, species-name & “pathogen”, species-name & “pig pathogen”. Publications were selected when describing early isolation of the bacterial species, or when describing pathogenicity (for example in a case study). We checked if the species had a safety status as assigned by the FDA (GRAS) or EFSA (QPS) (40) and we checked what risk group the species belonged to according to the German BAuA/ABAS’s Technical Rules for Biological Agents list (TRBA 466) (62). Species only belonging to TRBA risk group 1 were included in the species of interest list.

### Targeted isolation of bacteria

High relative abundances of the species of interest, in the 16S and *tuf* datasets, guided swab selection (top 5) for culturing (paired swabs were stored in 20% peptone-glycerol buffer frozen at -80°C). Per species of interest, we selected their suitable growth conditions (based on literature and web searches, Supplementary Table 1). The following media were selected: Brain Heart Infusion Agar (BHI), Columbia Agar with 5% Sheep Blood (Blood), Chocolate Agar (Choc), Glucose Yeast Peptone Agar (GYP), Luria-Bertani agar (LB), Marine Agar with 5% NaCl (MA5), De Man–Rogosa–Sharpe agar (MRS), Manitol Salt Agar (MSA), Nutrient Agar (NA), and Trypticase Soy Agar (TSA). Agar plates were ordered from commercial vendors except for NA, MRS, and MA5 which were prepared in house. Agar plates were incubated at either, 24°C, 30°C, 37°C, or 45°C, with one four gas mixtures (aerobic, aerobic +5% CO2 (20% O2, 5% CO2, 75% N2), anaerobic (0.2% O2, 10% CO2, 5% H2, 85% N2), and microaerobic (6% O2, 7% CO2, 3.6% H2, 84%N2), according to the expected need of the targeted species of interest. Gas mixtures were generated using the Anoxomat platform (Advanced Instruments, the Netherlands). Nasal swab samples were plated using 10µL loops. After overnight growth, plates were checked for growth. If no growth or tiny colonies were detected another 24 hours of growth was permitted. Upon colony formation, colonies with distinct morphology were picked with a 1 µL loop, re-streaked on fresh agar plates, and incubated for the same duration and under the same growth conditions as the original. Subculturing continued until the morphology of an isolate on agar appeared uniform and purity was assumed. Pure cultures were subsequently submitted for MALDI-TOF analysis.

### MALDI-TOF and 16S sequencing taxonomy assignment

Pure isolates were picked from the agar plate using a sterile toothpick and smeared onto the MALDI-TOF target *in duplo*. The smeared spots were treated with 1 µL 70% formic acid and left to dry. Dried spots were coated with 1 µL HCCA MALDI-TOF matrix (Bruker). Finished targets were analyzed on a MALDI-TOF MS platform (Bruker, Leiderdorp, the Netherlands). An isolate was considered as assigned genus taxonomy if the MALDI-TOF results were consistent between duplicates and had a score-value of >1.70, and species taxonomy was assigned at a score >2.00. Isolates that could not be assigned a genus or species were additionally 16S rRNA gene Sanger sequenced (Baseclear, Leiden, the Netherlands) using primers 27F (5’-AGA GTT TGA TYM TGG CTC AG-3’) and 1492R (5’TAC GGY TAC CTT GTT ACGACTT-3’). When genus of species identification matched the taxa of interest, colony-material was suspended in peptone-glycerol medium and preserved at -80°C.

### Tetracycline screen informed by EFSA guidance

A tetracycline resistance analysis was performed based on a per taxon defined tetracycline MIC-cutoff as guided by the 2018 EFSA guidance “Guidance on the characterization of microorganisms used as feed additives or as production organisms” (37). We prepared GYP and BHI agar plates containing tetracycline concentrations from 2 mg/L up till 16 mg/L with 2-fold increments. Isolate material was scraped from glycerol stocks and resuspended in 100 µL physiological saline solution. From this suspension isolates were spotted onto the selective plates and incubated at original isolate specific growth conditions. Plates were scored both after overnight growth and after another 24 hours.

### DNA extraction and whole genome sequencing

For DNA isolation bacteria were grown on agar plates. Bacterial colony material was re-suspended in 1 mL physiological salt solution with 1 mM EDTA. Material was spun at 10,000 RPM for 1 minute to pellet, and supernatant was removed. Cell pellets were used as input for the DNA extraction using the NGS DNeasy Ultraclean Microbial Kit (Qiagen Benelux BV, Venlo, the Netherlands) with bead beating according to manufacturer’s protocol. DNA sequencing libraries were prepared using the SQK-RBK110.96 rapid barcoding kit. Nanopore sequencing was conducted using a MinION with R9.4.1 flow cells (Oxford Nanopore Technologies). Base calling, barcode splitting, and barcode removal were performed using MinKNOW v5.4.3 with the “Super accurate” model. Reads were assembled to contigs with Flye v2.9 (2). Assemblies were polished using Medaka 1.4.3 (68) and Homopolish 0.3.4 (69). Species were determined directly from reads using WIMP in Epi2Me (https://epi2me.nanoporetech.com/), and using Kmerfinder (70–72) and TYGS (73) on assembled genomes. Resistance genes were scored using Resfinder (74) with a minimum coverage of 60% and an minimum identity of 80%.

### Phenotypic AMR profiling

Minimum inhibitory concentrations (MIC) were determined by broth microdilution using microtiter panels (ThermoFisher Scientific, Bleiswijk, The Netherlands and MERLIN diagnostika GmbH, Bonn, Germany). Additionally, we used clindamycin and streptomycin E-tests (Biomerieux Benelux, Amersfoort, The Netherlands) Summary of the ranges used can be found in Supplementary Table 5. Breakpoints from the 2018 EFSA guidance “Guidance on the characterization of microorganisms used as feed additives or as production organisms (Table 2)” (37) were used for interpretation of the results.

### *in vitro* efficacy testing of candidates against *S. aureus* strains

To assess *in vitro S. aureus* inhibiting capacity of the three selected *Lactococcus lactis* strains we performed a spent medium assay. To generate spent medium, strains were incubated for 20 hours in 100 mL BHI broth shaking at 37°C in a 250 mL Schott-bottle with the cap loose. After growth, we filtered cells at 0.45 µm and 0.20 µm pore size to generate the sterile spent medium. Spent medium was used to create four test conditions. Plain spent medium, spent medium with 50% fresh BHI medium added, spent medium with its pH adjusted to 7.4, and spent medium with 50% BHI and pH adjustment to 7.4. As a positive control plain BHI was used from the same batch used for the spent medium. 200 µL of the treatment reaction mixtures was placed in a 96 wells plate and inoculated with 1 µL of cell suspension of the test strain at OD_600nm_ 0.6-0.8. Plates were covered and incubated shaking overnight at 37°C in a plate reader (Synergy HTX, Agilent BioTek) that measured OD_600 nm_ at 10-minute intervals.

## Figures and tables

**Supplementary Table 1.** Overview of collected information of all the anticorrelating species and their selection (of interest column).

**Supplementary Table 2.** Grown isolates with taxonomy assignment and their information.

**Supplementary Table 3.** An overview of all performed assays on the isolates of interest.

**Supplementary Table 4.** pH of the liquid culture media before and after pH correction

**Supplementary Table 5.** Antibiotic range tested.

## Author contributions

AV, JW, BD, AZ, PL, MC designed the study. DCP, AV, JW and CE collected the pig samples. JE, CH, and AV carried out the amplicon sequencing. SP, AV, and AZ performed the data analysis. AV, MC, BD, AZ, JW, and PL participated in the interpretation of the results. AV wrote the manuscript with input from all other authors. All authors read and approved the final manuscript.

## Conflicts of interest

The author(s) declare that there are no conflicts of interest.

## Funding information

This research was funded by The Irish Health Research Board (ExcludeMRSA, grant number JPI-AMR-2017-1-A) and in part by Science Foundation Ireland (grant numbers SFI/12/RC/2273_P2 and 17/CDA/4765). The Dutch research was funded by ZonMw (The Netherlands Organisation for Health Research and Development), project number 541002003, and JPIAMR (JPIAMR-2017-1-B grant number 50-52900-98-043).

## Consent for publication

Informed consent by the participating farmers was obtained.

## Ethical approval

Approval for materials collected were described previously in Vlasblom *et al.*, 2024 (https://www.biorxiv.org/content/10.1101/2023.12.20.572551v1).

For current work no approval was required.

## Availability of data and materials

STORMS files, scripts, metadata including qPCR results, and Phyloseq objects are available through ZENODO: https://doi.org/10.1101/2023.12.20.572551

Sequencing reads of *tuf* and 16S rDNA are available under accession number: PRJEB71383 https://www.ebi.ac.uk/ena/browser/view/PRJEB71383).

Strain whole genome reads are available under accession number: PRJEB76400

## Acknowledgements

We want to thank Marian Broekhuizen-Stins, Maximilian Casteel, Gerard van Eijden, Demi van Dijk, Demi van de Hoef, Arjen Timmerman, and Heleen Zweerus, for their contributions to the sampling, and sample-processing effort.

